# Improved bacterial recombineering by parallelized protein discovery

**DOI:** 10.1101/2020.01.14.906594

**Authors:** Timothy M. Wannier, Akos Nyerges, Helene M. Kuchwara, Márton Czikkely, Dávid Balogh, Gabriel T. Filsinger, Nathaniel C. Borders, Christopher J. Gregg, Marc J. Lajoie, Xavier Rios, Csaba Pál, George M. Church

## Abstract

Exploiting bacteriophage-derived homologous recombination processes has enabled precise, multiplex editing of microbial genomes and the construction of billions of customized genetic variants in a single day. The techniques that enable this, Multiplex Automated Genome Engineering (MAGE) and directed evolution with random genomic mutations (DIvERGE), are however currently limited to a handful of microorganisms for which single-stranded DNA-annealing proteins (SSAPs) that promote efficient recombineering have been identified. Thus, to enable genome-scale engineering in new hosts, highly efficient SSAPs must first be found. Here we introduce a high-throughput method for SSAP discovery that we call Serial Enrichment for Efficient Recombineering (SEER). By performing SEER in *E. coli* to screen hundreds of putative SSAPs, we identify highly active variants PapRecT and CspRecT. CspRecT increases the efficiency of single-locus editing to as high as 50% and improves multiplex editing by 5 to 10-fold in *E. coli*, while PapRecT enables efficient recombineering in *Pseudomonas aeruginosa*, a concerning human pathogen. CspRecT and PapRecT are also active in other, clinically and biotechnologically relevant enterobacteria. We envision that the deployment of SEER in new species will pave the way toward pooled interrogation of genotype-to-phenotype relationships in previously intractable bacteria.

## Introduction

The increasing accessibility of whole-genome sequencing to microbiologists has created a gap between researchers’ ability to read vs. their ability to write/edit genetic information. As the availability of sequencing data increases, so too does its hypothesis-generating capacity, which motivates the development of genome-editing tools that can be employed to create user-defined genetic variants in a massively parallelizable manner. Currently, most techniques for editing microbial genomes that meet these criteria exploit bacteriophage-derived homologous recombination processes^1–3^. Molecular tools derived from phages enabled recombineering (recombination-mediated genetic engineering), which uses short homology arms to efficiently direct double-stranded DNA (dsDNA) cassette- and single-stranded DNA (ssDNA)-integration into bacterial genomes^4–6^. Improvements to recombineering in *E. coli* enabled multiplex, genome-scale editing and the construction of billions of genetic variants in a single experiment^7–9^. Two of these recombineering-based methods, Multiplex Automated Genome Engineering (MAGE) and directed evolution with random genomic mutations (DIvERGE) have been used for a variety of high-value applications^3, 10–14^, but at their core they offers the ability to generate populations of bacteria that can contain billions of precisely targeted mutations. However, while these techniques function well in *E. coli* and some closely related Enterobacteria, efforts to reproduce these results in other bacterial species have been sporadic and stymied by low efficiencies (Table S1)^15–31^.

The incorporation of genomic modifications via oligonucleotide annealing at the replication fork, called oligo-mediated recombineering, is the molecular mechanism that drives MAGE and DIvERGE^4, 32, 33^. This method was first described in *E. coli*, and is most commonly promoted by the expression of *bet* (here referred to as Redβ) from the Red operon of Escherichia phage λ^5, 6, 34^. Redβ is a single-stranded DNA-annealing protein (SSAP) whose role in recombineering is to anneal ssDNA to complimentary genomic DNA at the replication fork. Though improvements to recombineering efficiency have been made^7–9^, the core protein machinery has remained constant, with Redβ representing the state-of-the-art in *E. coli* and enterobacterial recombineering. Redβ additionally does not adapt well to use outside of *E. coli*, displaying host tropism, presumably toward hosts that are targets of infection for Escherichia phage λ. To enable recombineering in organisms in which Redβ does not work efficiently, most often Redβ homologs from prophages of genetically similar bacteria are screened^15, 16, 21, 23, 30^, but high levels of recombineering efficiency, as seen in *E. coli*, remain elusive.

We hypothesized that the identification of an optimal SSAP is currently the limiting factor to improved recombineering efficiency and to developing multiplex genome engineering tools (i.e. MAGE) in new bacterial species. To provide a solution, here we present a high-throughput method for isolating SSAP homologs that efficiently promote recombineering, which we call Serial Enrichment for Efficient Recombineering (SEER) (Figure 1). SEER allowed us to screen two SSAP libraries in a matter of weeks, each of which contains more members than, to our knowledge, the sum total from all previous SSAP-screening efforts (Table S1). We first use SEER to screen a library of 122 SSAPs from seven different families: RecT, Erf, Sak4, Gp2.5, Sak, Rad52, and RecA, finding that proteins of the RecT and Erf families are the most promising candidates for future screening. In a follow-on screen of 107 RecT variants, we then identify CspRecT (from a *<u>C</u>ollinsella <u>s</u>tercoris* <u>p</u>hage), which doubles recombineering efficiency in *E. coli* over Redβ. Next, by focusing on another enriched variant, PapRecT (from a *<u>P</u>seudomonas <u>a</u>eruginosa* <u>p</u>hage), we demonstrate high efficiency genome editing in its native host *Pseudomonas aeruginosa*, which has long lacked good genetic tools. We then broadly profile the activity of CspRecT and PapRecT across diverse Gammaproteobacteria. Following on our successful demonstration in this work, we believe that SEER should be an easily adaptable method for identifying optimal proteins for recombineering across a wide range of bacterial species.

**Figure 1.**
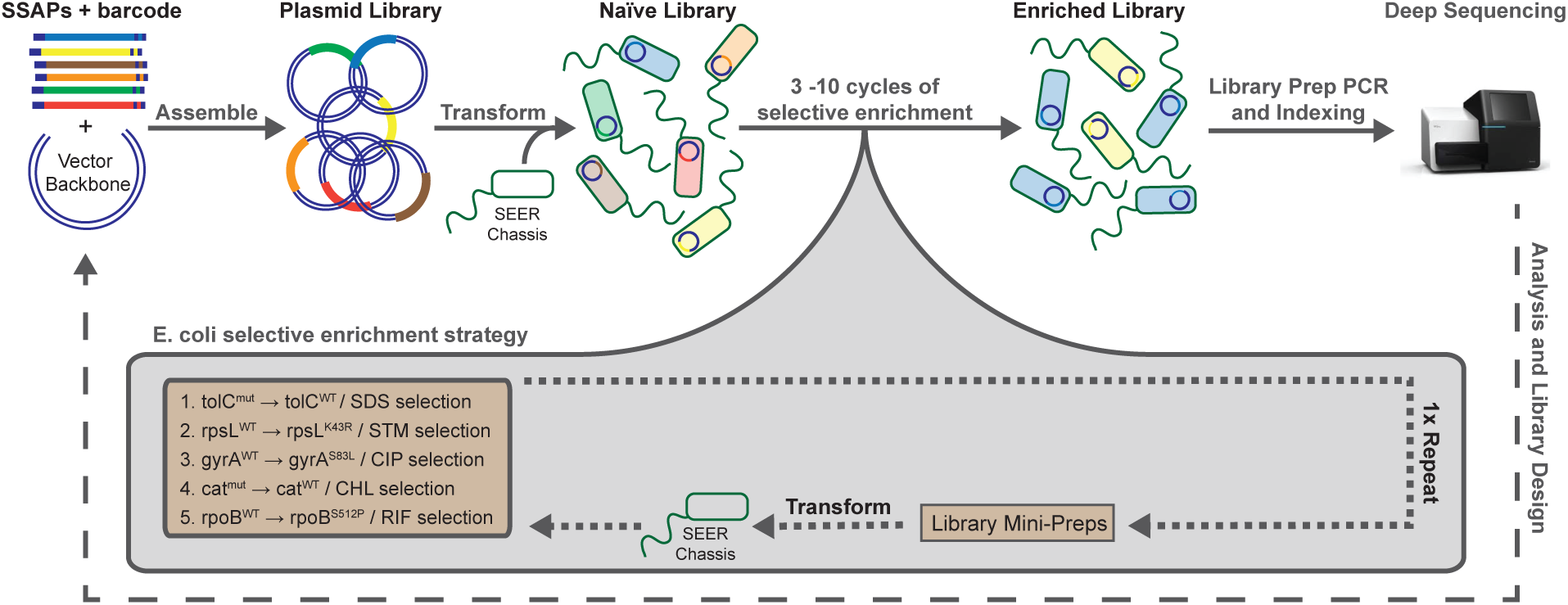
Serial Evolutionary Enhancement of Recombineering (SEER) workflow. The SEER workflow is depicted across the top from left to right, with libraries being first assembled, then moved into a chassis organism, enriched over 3-10 cycles for efficient recombineering proteins, and finally analyzed by deep sequencing. This process can then be iterated on by learning from results and making improvements to the library design. The specific selective enrichment strategy that we designed for *E. coli* is shown in a gray callout bubble. Five successive antibiotic selections were applied to a library of *E. coli* cells expressing SSAP variants, and after the selective handles were exhausted the plasmid library was extracted and re-transformed into the naïve SEER chassis for five further cycles of selection.

## Results and Discussion

### Identification of single-stranded DNA-annealing proteins (SSAPs)

Previous analyses of phylogenetic data suggest that there are seven principal families of phage-derived SSAPs: RecT (Pfam family: PF03837), Erf (Pfam family: PF04404), Rad52 (Pfam family: PF04098), Sak4 (Pfam family: PF13479), Gp2.5 (Pfam family: PF10991), Sak (Pfam family: PF06378), and RecA (Pfam family: PF00154)^18, 35^. These have been further grouped into three superfamilies, wherein RecT, Erf, and Sak have been proposed to adopt Rad52-like folds and Sak4 and RecA cluster together into a Rad51-like superfamily^18^. Furthermore, Redβ homologs are best classified as a part of the larger RecT family, and UvsX homologs are best classified in the RecA protein family^36^. To generate a library that widely sampled SSAP diversity, we used a Hidden Markov Model to search metagenomic databases beginning with an ensemble of SSAPs that have demonstrated activity in *E. coli*^19, 37^ (see methods). A library of 131 proteins was identified and codon-optimized for expression in *E. coli*, 121 of which were synthesized without error (Table S2) and cloned into a plasmid vector with a standard arabinose-inducible expression system. This assembled group of SSAP variants, which we refer to here as *Broad SSAP Library*, has members from all seven families of SSAPs (Figure 2A) and is phylogenetically diverse (Figure 2B). To facilitate tracking the library over multiple selective cycles, we added 12-nucleotide barcode 22 nucleotides downstream of the stop codon of each gene (Figure S1), which enabled us to identify each SSAP variant by PCR-amplification and targeted Next Generation Sequencing (NGS).

**Figure 2.**
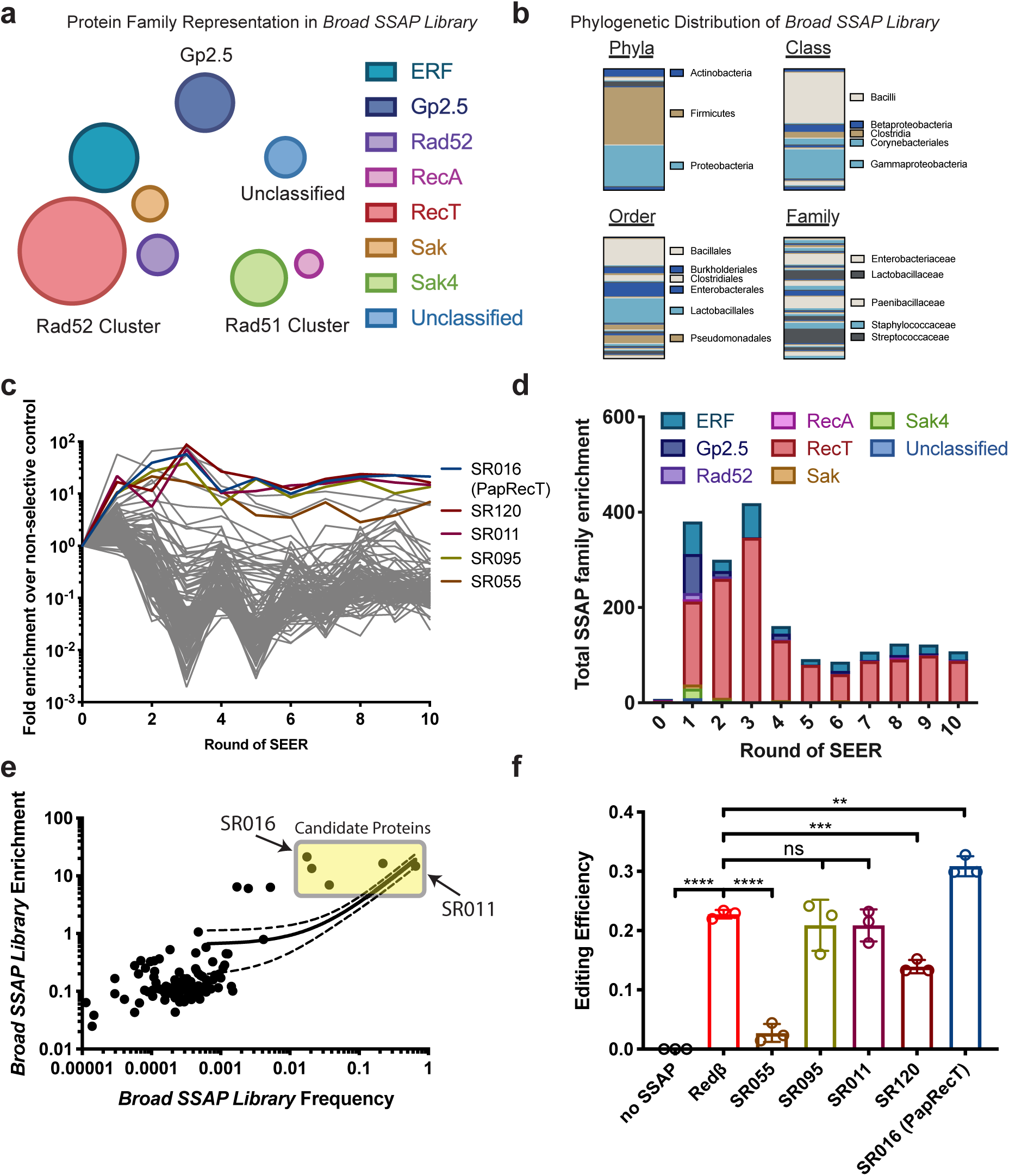
*Broad SSAP Library* (A) Circles of various sizes represent the number of variants present in *Broad SSAP Library* from different protein families, as categorized by the Pfam database. The seven principle families of SSAPS are grouped into three clusters based on structural and phylogenetic information as proposed by Lopes et al. (2010). (B) The phylogenetic distribution of *Broad SSAP Library* is represented as vertical bar charts. For each phylogenetic level, any group that represents more than 5% of the total library is called out to the right of the bar. (C) The enrichment of each *Broad SSAP Library* member is plotted over ten successive rounds of selection. Enrichment of each library member was calculated by dividing the average frequency across selective replicates by the frequency of the non-selective control. Frequency data is measured by amplification of the barcoded region of the plasmid library and next generation sequencing (NGS). (D) Total enrichment is plotted for each protein family over the ten rounds of SEER. (E) Frequency is plotted against enrichment for each *Broad SSAP Library* member after the tenth round of selection. Five candidate proteins, which are shaded in a yellow box, were selected for further characterization. (F) Five top candidates were tested for their efficiency at incorporating a single-base-pair silent mutation at a non-essential gene, *ynfF*. Efficiency was read out by NGS. Significance values are indicated for a parametric t-test between two groups, where ns, *, **, ***, and ***** indicate p > 0.05, p < 0.05, p < 0.01, p < 0.001, and p < 0.0001 respectively.

### Serial Enrichment for Efficient Recombineering (SEER)

Next, to evaluate libraries of computationally identified SSAP variants, we established a high-throughput assay to select for SSAPs that promote efficient recombineering. We hypothesized that the iterative, serial integration of easily selectable mutations into the bacterial genome, coupled to subsequent allelic selection, could allow us to select SSAP variants *en masse*, and thus enrich best performers from diverse SSAP libraries (Figure 1). This method, that we term Serial Enrichment for Efficient Recombineering (or SEER) proceeds via i.) the identification and cloning of SSAP libraries into expression vectors, ii.) the transformation of such libraries into an organism-of-interest, followed by iii.) enrichment for SSAPs efficient at recombineering, and then iv.) analysis by deep sequencing to read out library composition. Step “iii” comprises the successive transformation of oligos, which upon successful integration into the host genome will confer a resistant phenotype, followed by antibiotic selection for the resistant allele across the entire library in liquid culture. SSAP variants that incorporate mutations most effectively will thereby be enriched. Finally, if needed, after the selective handles are exhausted, the SSAP-plasmid library can be extracted and re-transformed into a naïve chassis for further cycles of enrichment.

To perform the selections efficiently we engineered an *E. coli* chassis to both improve recombineering efficiency and to introduce genetic handles with which we could apply selective pressure against non-edited alleles. We used as our parental strain, EcNR2^3^, a derivative of *E. coli* K-12 MG1655 that has its methyl-directed mismatch repair (MMR) machinery disabled (Δ*mutS*::*cat*) to improve recombineering efficiency (see Methods for details)^7^. EcNR2 was modified to i.) improve recombineering efficiency (DnaG Q576A)^8^ and to ii.) introduce genetic handles for antibiotic selection. To this latter aim, stop codons were introduced into both *cat* (in the *mutS* locus) and *tolC*, making the modified strain sensitive to chloramphenicol (CHL) and sodium dodecyl sulfate (SDS). We refer to this modified organism as the “SEER chassis”. Next, to identify selectable markers beyond *cat* and *tolC*, and thus allow multiple SEER selection cycles, we tested a group of resistant alleles that we gathered from an analysis of antibiotic resistance literature (See Methods for details). We found GyrA_S83L to confer robust resistance to ciprofloxacin (CIP), RpoB_S512P to rifampicin (RIF), and RpsL_K43R to streptomycin (STR) (Figure S2). We then verified that these three antibiotic selections, in addition to the CHL and SDS selections described above, are orthogonal and that they could be performed serially on our engineered *E. coli* SEER chassis. This enabled us to run ten rounds of SEER with only a single extraction and re-transformation of the plasmid libraries into the naïve chassis (Figure 1).

Following the optimization of the SEER workflow, we performed ten successive cycles of selection on *Broad SSAP Library*, representing all major SSAP families, in the *E. coli* SEER chassis. Four replicate populations were run: one control population underwent induction of protein expression and oligo transformation, but without antibiotic selection, while three experimental replicates underwent antibiotic selections in the following order: SDS → STR → CIP → CHL → RIF (Figure 1C). To track the enrichment of each library member we amplified the 12-nucleotide barcode region after each round of selection and sequenced by NGS (Table S3). Clear winners emerged relatively quickly, with top variants increasing in frequency distinctly after only a few rounds of selection (Figure S3). The non-selected control population, however, also displayed significant enrichment effects, which indicated that the over-expression of certain SSAP variants may impair fitness. Thus, to normalize for fitness and the cost of protein expression, we next calculated an enrichment score for each library member: we divided the average frequency of each variant within the selected populations by its frequency in the non-selected control population (Figure 2C). Based on NGS data, we found that the RecT family was the most enriched family by a large margin (85-fold over the non-selected control), followed by the ERF family (18-fold over the non-selected control), and no other SSAP family showed significant enrichment (Figure 2D). However, it is important to note that our codon-optimized Redβ (SR085 in our library) was not significantly enriched through the ten rounds of selection. To investigate this issue further we compared the performance of Redβ expressed off of its wild-type codons against the codon-optimized version that was included in *Broad SSAP Library*. This revealed significantly decreased efficiency for the codon-optimized version of Redβ (Figure S4), which indicates that codon choice is an important consideration for library design, and that negative results for individual SSAPs from a large screen of this sort do not necessarily indicate that the protein itself is not functional.

Finally, we chose a set of five library members that exhibited both high frequency and enrichment for further analysis (Figure 2E). We tested their recombineering efficiency against Redβ expressed off of its wild-type codons on the same plasmid system used for the SEER selections. To ensure an accurate measurement we queried the efficiency of each SSAP by NGS after performing a silent, non-coding genetic mutation at a non-essential gene, *ynfF* (silent mismatch MAGE oligo 7: Table S4). *Broad SSAP Library* member: SR016, which we introduced earlier as PapRecT (UniParc ID: UPI0001E9E6CB), demonstrated the highest efficiency of recombineering, i.e. 31% ± 2% (Figure 2F).

### Screening diverse RecT homologs identifies a highly efficient SSAP

By investigating the distribution of efficient recombineering-functions across the seven principal families of phage-derived SSAPs, our first SEER screen suggested the RecT family (Pfam family: PF03837) as the most abundant source of recombineering proteins for *E. coli*. Importantly, previous screens by others also detected efficient SSAPs from the RecT family (Table S1). Therefore, we hypothesized that by screening additional RecT variants, again exploiting the increased throughput of SEER compared to previous efforts, we might discover recombineering proteins further improved over Redβ and PapRecT. To this aim we constructed a second library, identifying a maximally diverse group of 109 RecT variants, 106 of which were synthesized successfully, which we call *Broad RecT Library* (see Methods for more details). Next, as previously described, we performed 10 rounds of SEER selection on *Broad RecT Library* (Figure S5), and upon plotting frequency against enrichment after the final selection, a clear winner emerged (Figure 3a). This protein, which we introduced earlier as CspRecT (UniParc ID: UPI0001837D7F), originates from a phage of the Gram-positive bacterium *Collinsella stercoris*.

**Figure 3.**
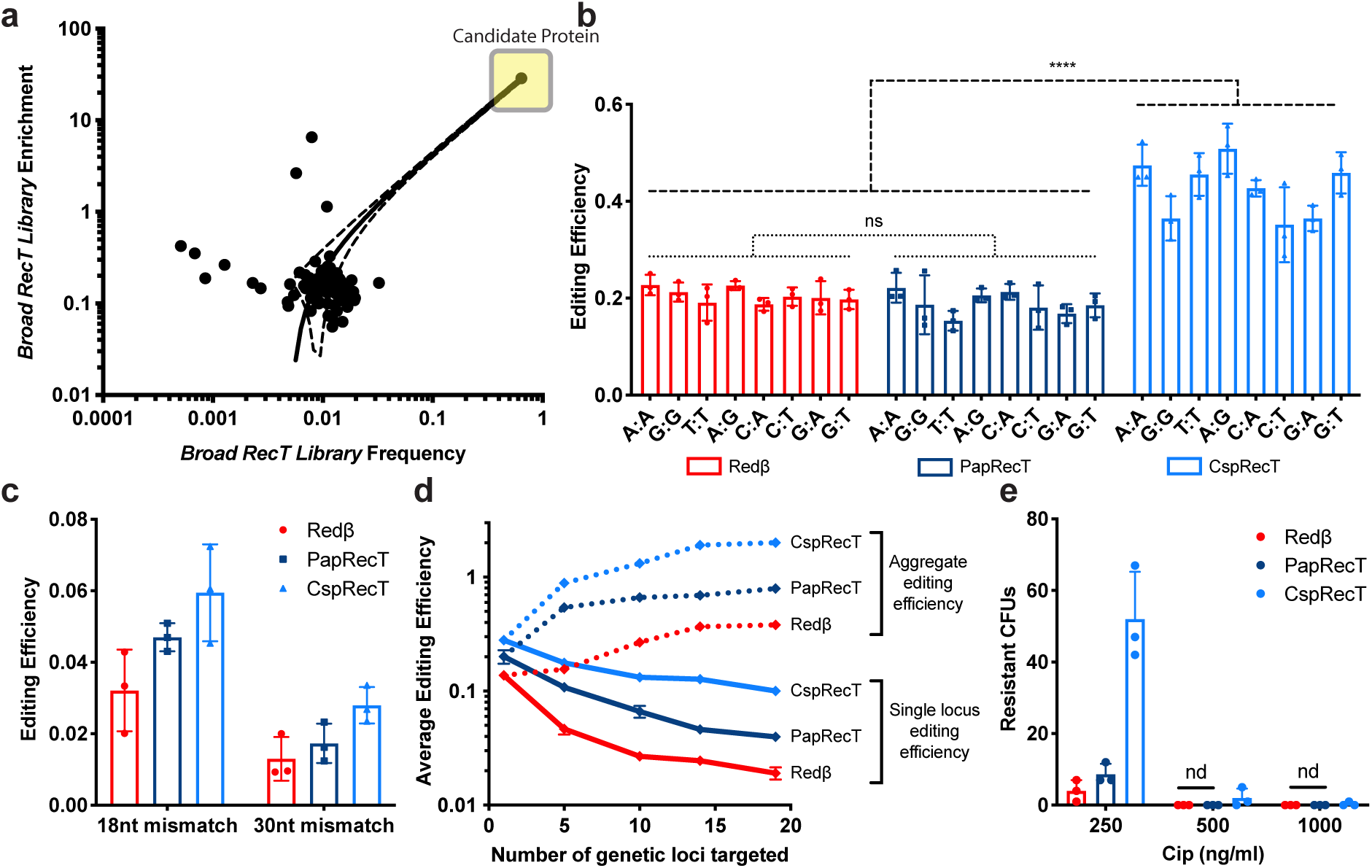
*Broad RecT Library* and CspRecT (A) Frequency is plotted against enrichment for each *Broad RecT Library* member after the tenth round of selection. One candidate protein, CspRecT (shaded box), was the standout winner. In all subsequent panels, Redβ, PapRecT, and CspRecT are compared when expressed from a pORTMAGE-based construct (Figure S1) in wild-type MG1655 *E. coli*. Significance values are indicated for a grouped parametric t-test, where ns and ***** indicate p > 0.05 and p < 0.0001 respectively. Editing efficiency was measured by blue/white screening at the LacZ locus for (B) eight different single-base mismatches (n=3) and (C) 18-base and 30-base mismatches (n=3). (D) MAGE editing targeting 1, 5, 10, 15, or 20 genomic loci at once in triplicate, was read out by NGS. The solid lines represent the mean editing efficiency across all targeted loci, while the dashed lines represent the sum of all single-locus efficiencies, which we refer to as aggregate efficiency. (E) A 130-oligo DIvERGE experiment using oligos that were designed to tile four different genomic loci that encode the drug targets of fluoroquinolone antibiotics and are known hotspots for CIP resistance. The oligos contained 1.5% degeneracy at each nucleotide position along their entire length. All 130 oligos were mixed and transformed together into cells (n=3). Colony forming units were measured at three different CIP concentrations after plating 1/100^th^ of the final recovery volume.

To maximize the phylogenetic reach and applicability of these new tools, we characterized CspRecT, alongside Redβ and PapRecT, subcloned into the pORTMAGE plasmid system (Figure S1, Addgene accession: 120418). This plasmid contains a broad-host RSF1010 origin of replication, establishes tight regulation of protein expression with an m-toluic-acid inducible expression system, and disables MMR by transient overexpression of a dominant-negative mutant of *E. coli* MutL (MutL E32K)^38^, which makes it possible to establish high-efficiency editing without modification of the host genome. Measured with a standard lacZ recombineering assay, wild-type *E. coli* MG1655 expressing CspRecT exhibited editing efficiency of 35-51% for various single-base mismatches, averaging 43% or more than double the efficiency of cells expressing Redβ or PapRecT off of the same plasmid system (Figure 3B). This pORTMAGE plasmid expressing CspRecT we refer to as pORTMAGE-Ec1 (Addgene accession: 138474). To our knowledge the efficiency of CspRecT single-locus genome editing reported here is the first to significantly exceed 25%, the theoretical maximum for a single incorporation event^39^, implying that editing occurs either at multiple forks or over successive rounds of genome replication.

We then tested CspRecT at a variety of more complex genome editing tasks. For longer strings of consecutive mismatches, which are lower efficiency events, CspRecT was again about twice as efficient as Redβ. Wild type *E. coli* MG1655 expressing CspRecT displayed 6% or 3% efficiency (vs. 3% or 1% for Redβ) for the insertion of oligos conferring 18-bp or 30-bp consecutive mismatches into the lacZ locus respectively (Figure 3C). To further investigate the performance of CspRecT at complex, highly multiplexed genome editing tasks, we designed a set of 20 oligos spaced evenly around the *E. coli* genome, each of which incorporates a single-nucleotide synonymous mutation at a non-essential gene. Next, while expressing Redβ, PapRecT, and CspRecT separately from the corresponding pORTMAGE plasmid in *E. coli* MG1655, we performed a single cycle of genome editing with equimolar pools of 1, 5, 10, 15, and 20 oligos and assayed editing efficiency at each locus by PCR amplification coupled to targeted next generation sequencing (NGS). NGS analysis revealed a general trend: as the number of parallel edits grew, the degree of overperformance by CspRecT also grew (Figure 3D). For instance, when making 19 simultaneous edits (one oligo from the pool of 20 could not be read out due to inconsistencies in allelic amplification), CspRecT averaged 10.0% editing efficiency at all loci, whereas PapRecT averaged 4.0% and Redβ averaged 1.9%. Importantly, despite keeping total oligo concentration fixed across all pools, aggregate editing efficiency increased as more oligos were present in each pool. For instance, when using CspRecT with a 19-oligo pool, aggregate editing efficiency was more than 200%, implying that across the total recovered population of *E. coli* there averaged more than two edits per cell.

Finally, based on the increased integration efficiency with CspRecT in multiplexed genome editing tasks, we also tested its performance in a directed evolution with random genomic mutations (DIvERGE) experiment^13^. DIvERGE uses large libraries of soft-randomized oligos that have a low basal error rate at each nucleotide position along their entire sequence to incorporate mutational diversity into a targeted genomic locus. To compare the performance of Redβ, PapRecT, and CspRecT, we performed one round of DIvERGE mutagenesis by simultaneously delivering 130 partially overlapping DIvERGE oligos designed to randomize all four protein subunits of the drug targets of ciprofloxacin (*gyrA*, *gyrB*, *parC*, and *parE*) in *E. coli* MG1655. Following library generation, cells were subjected to 250, 500, and 1,000 ng/mL ciprofloxacin (CIP) on LB-agar plates. Variant libraries that were generated by expressing CspRecT produced more than ten times as many colonies at low CIP concentrations (i.e., 250 ng/mL) as Redβ and PapRecT, while at 1,000 ng/mL CIP, which requires the simultaneous acquisition of at least two mutations (usually at *gyrA* and *parC*) to confer a resistant phenotype, only the use of CspRecT produced resistant variants (Figure 3E). Because *gyrA* and *parC* mutations are usually necessary to confer high-level CIP resistance, sequence analysis of *gyrA* and *parC* from 11 randomly selected CIP-resistant colonies, we found many different mutations, in combinations of up to three, most of which have been described in resistant clinical isolates^13, 40^ (Table S5). In sum, in both MAGE and DIvERGE experiments, which require multiplex editing, CspRecT provided more than an order of magnitude improvement to editing efficiency over Redβ, the current state-of-the-art recombineering tool.

### Improved genome editing in diverse Gammaproteobacteria

SSAPs frequently show host tropism^15, 25, 26^, but there are also indications that within bacterial clades certain SSAPs may function broadly^21, 38, 41^. Therefore, we next investigated the functionality of PapRecT and CspRecT in selected Gammaproteobacteria and compared their efficiency to that of Redβ. We chose to focus our efforts on two enterobacterial species: *Citrobacter freundii* ATCC 8090 and *Klebsiella pneumoniae* ATCC 10031, along with the more distantly related *Pseudomonas aeruginosa* PAO1. Pathogenic isolates of *K. pneumoniae* and *P. aeruginosa* are among the most concerning clinical threats due to widespread multidrug resistance^42^. In these species, oligo-recombineering based multiplexed genome editing (i.e., MAGE and DIvERGE) holds the promise of enabling rapid analysis of genotype-to-phenotype relationships and predicting future mechanisms of antimicrobial resistance^13, 43^. *C. freundii*, by contrast, is an intriguing biomanufacturing host in which the optimization of metabolic pathways has remained challenging^44, 45^

To test the activity of PapRecT and CspRecT in these three organisms we built on the broad-host-range pORTMAGE system^38^ described above. For experiments in *E. coli* we had subcloned PapRecT or CspRecT in place of Redβ into pORTMAGE311B^46^ (Figure S1; Addgene accession: 120418), which transiently disrupts MMR with *Ec*MutL_E32K, and whose RSF1010 origin of replication and m-toluic-acid-based expression system allows the plasmid to be deployed over a broad range of bacterial hosts^47, 48^. We used these same pORTMAGE-based constructs for testing in *C. freundii* and *K. pneumoniae*. In *P. aeruginosa* the plasmid architecture remained constant, except that we replaced the origin of replication and antibiotic resistance, instead using the broad-host-range pBBR1 origin, which was shown to replicate in *P. aeruginosa*^49^, and a gentamicin resistance marker (Figure S1). Next we tested these constructs (See methods for details), and in all three species, PapRecT and CspRecT displayed high editing efficiencies (Figure 4A). In *C. freundii* and *K. pneumoniae*, just as in *E. coli*, we found CspRecT to be the optimal choice of protein, whereas in *P. aeruginosa* PapRecT performed the best. We further compared PapRecT to two recently reported *Pseudomonas putida* SSAPs (Rec2 and Ssr)^26, 50^, and found that PapRecT, isolated from a large *E. coli* screen performed equal to or better than proteins found in smaller screens run through *P. putida* (Figure S6). We found, however, that the efficiency of our plasmid construct was lower in *P. aeruginosa* than in the enterobacterial species that pORTMAGE was optimized for. Therefore, to increase editing efficiency in *P. aeruginosa*, we next i.) optimized ribosomal binding sites (RBS) for PapRecT and *Ec*MutL, ii.) replaced *Ec*MutL_E32K with its equivalent homologous mutant from *P. aeruginosa* (*Pa*MutL_E36K), iii.) incorporated the native *P. aeruginosa* coding sequence for PapRecT instead of the *E. coli* codon-optimized version (Figure S7). Together these changes significantly improved the editing efficiency of our best plasmid construct featuring PapRecT in *P. aeruginosa*, which we call pORTMAGE-Pa1 (Addgene accession: 138475), to ∼15%.

**Figure 4.**
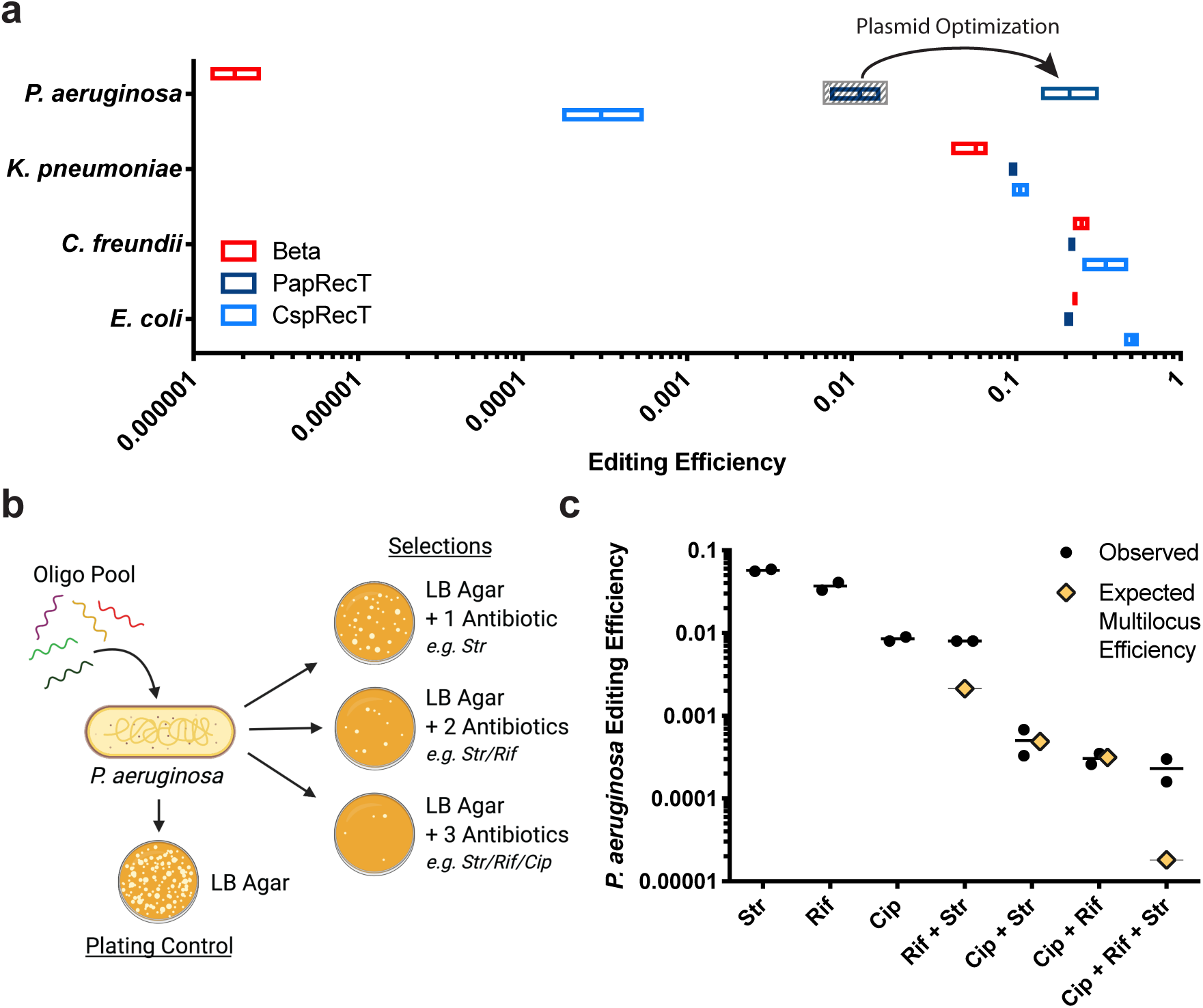
Recombineering in Gammaproteobacteria. (A) Recombineering experiments were run with Redβ, PapRecT, and CspRecT expressed off of the pORTMAGE311B backbone, or with a pBBR1 origin in the case of *P. aeruginosa*. Editing efficiency was measured by colony counts on selective vs. non-selective plates (n=3; see methods). Vector optimization resulted in improved efficiency of PapRecT in *P. aeruginosa* (see Figure S7) (B) Diagram of a simple multi-drug resistance experiment in *P. aeruginosa* harboring an optimized PapRecT plasmid expression system, pORTMAGE-Pa1. In a single round of MAGE, a pool of five oligos was used to incorporate genetic modifications that would provide resistance to STR, RIF, and CIP (n=3). These populations were then selected by plating on all combinations of 1-, 2-, or 3-antibiotic agarose plates and compared with a non-selective control. (C) Observed efficiencies were calculated by comparing colony counts on selective vs. non-selective plates. Expected efficiencies for multi-locus events were calculated as the product of all relevant single-locus efficiencies.

Virulent strains of *P. aeruginosa* are a frequent cause of acute infections in healthy individuals, as well as chronic infections in high-risk patients, such as those suffering from cystic fibrosis^51^. The rate of antibiotic resistance in this species is growing, with strains adapting quickly to all clinically applied antibiotics^52, 53^. The development of multidrug resistance in *P. aeruginosa* requires the successive acquisition of multiple mutations, but due to the lack of efficient tools for multiplex genome engineering in *P. aeruginosa*^54, 55^, investigation of these evolutionary trajectories has remained cumbersome. Therefore, and to demonstrate the utility of pORTMAGE-Pa1-based MAGE in *P. aeruginosa*, we simultaneously incorporated a panel of genomic mutations that individually confer resistance to STR, RIF, and fluoroquinolones (i.e., CIP)^56, 57^. Importantly, the corresponding genes are also clinical antibiotic targets in *P. aeruginosa*^58^. Following a single cycle of MAGE delivering 5 mutation-carrying oligos, a single-day experiment with pORTMAGE-Pa1, we were able to isolate all possible combinations of five resistant mutations, with more than 10^5^ cells from a 1ml overnight recovery attaining simultaneous resistance to STR, RIF, and CIP (Figure 4B). Interestingly, because *rpsL* and *rpoB*, the resistant loci for STR and RIF respectively, are located only ∼5kb apart from each other on the *P. aeruginosa* genome, these two mutations co-segregated much more often than would be expected by independent inheritance, confirming that co-selection functions similarly in *P. aeruginosa* to *E. coli* (Figure 4C)^12^. By genotyping and characterizing resistant colonies, we could then easily determine the Minimum Inhibitory Concentration (MIC) of CIP for various resistant genotypes (Table 1, Figure S8). The allure of this method is that the entire workflow took only three days to complete, in contrast with other genome engineering methods (i.e., CRISPR/Cas9 or base-editor-based strategies) that are either less effective, have biased mutational spectra, and/or would require tedious plasmid cloning and cell manipulation steps^54, 55^.

**Table 1.** Minimum inhibitory concentration (MIC) was measured for various combinations of ciprofloxacin resistance-conferring alleles. GyrA_T83I displays strong positive epistasis with ParC_S87L, and so clonal populations with mutations to *parC* but not *gyrA* were not pulled out of our antibiotic selection^75^.

| Genotype | Ciprofloxacin MIC (µg/ml) |
| --- | --- |
| PAO1 wild-type | 0.25 |
| nfxB knockout | 4 |
| GyrA_T83I | 16 |
| GyrA_T83I + ParC_S87L | 32 |
| GyrA_T83I + ParC_S87L + NfxB knockout | >128 |

## Conclusion

The discovery and subsequent improvement of recombineering in *E. coli* enabled the development of advanced genome-engineering techniques such as MAGE^3^, TRMR^59^, REXER^1^, Retron-Recombineering^10, 60^, and DIvERGE^13^, and applications ranging from genomic recoding^61^, biocontainment^11^, and viral resistance^61^ to the study and prediction of antibiotic resistance^13^ and the improved bioproduction of chemicals^12^. These powerful techniques all rely on efficient recombineering, but despite the essential role that an optimal SSAP plays, no method had hitherto been developed to rapidly and efficiently screen complex libraries of SSAPs. Here we describe such a method that we term Serial Enrichment for Efficient Recombineering (SEER), a technique for isolating efficient SSAPs from a large number of *in silico*-identified candidates, in which successive rounds of recombineering are followed by allelic selection (Figure 1). In the current work we have used SEER to screen two large libraries of SSAPs through *E. coli* and demonstrated the power of this method by isolating a variant, CspRecT that doubles the already high single-locus efficiency in *E. coli* to around 50%, a standout number and the highest reported gene-editing efficiency of any system that is not nuclease-dependent that we are aware of. We also demonstrate that CspRecT radically improves the multiplexability of oligo recombineering in *E. coli*, showing 5 to 10-fold improvement over Redβ with methods that rely on multiplex editing such as MAGE and DIvERGE.

Beyond the success of SEER in *E. coli*, the true promise of the technique is to increase the number of bacterial species in which these powerful genome-engineering tools are available to researchers. To this end we demonstrate that SEER can discover recombineering proteins that work not only in the targeted organism, but in closely related species as well. We show that proteins enriched in two SSAP libraries that were screened through *E. coli* meet or exceed the efficiency of the best tools available in *C. freundii*, *K. pneumoniae*, and *P. aeruginosa*. We report single-locus editing efficiencies of over 10% in each of these clinically and biotechnologically relevant Gammaproteobacteria, which should readily enable multiplex genome engineering.

However, it is also important to highlight the current limitations to the scope of this work. To expand modern genome-engineering tools to bacteria that are only distantly related to *E. coli*, SEER will need to be performed in new host organisms in the future. To this end we detail three important considerations for constructing SSAP libraries for future studies: i.) based on our screens in *E. coli*, and in concert with previous efforts (Table S1), the recT and ERF families are the most likely sources of SSAPs to drive efficient recombineering, ii.) phylogenetic diversity is essential, as CspRecT, the best-performing SSAP in *E. coli*, originates from a phage of *Collinsella stercoris*, a Gram-positive coriobacterium that is phylogenetically distant to *E. coli*, and iii.) future SSAP screening efforts should give careful consideration to codon optimization, as codon choice can have a large impact on apparent recombineering efficiency (as we observed with both Redβ and PapRecT). In short, SEER holds the promise of being a universal screening method that allows for the identification of highly active SSAPs in virtually any target bacterium, with the potential to broaden the applicability of recombineering-based genome-engineering tools toward as yet genetically intractable microorganisms.

## Materials and Methods

### Strains and Reagents

All strains used in this study are listed in Table S6. Unless otherwise noted, bacterial cultures were grown in Lysogeny-Broth-Lennox (LB^L^) (10g tryptone, 5g yeast extract, 5g NaCl in 1L H_2_O). Super optimal broth with catabolite repression (SOC) was used for recovery after electroporation. MacConkey agar (17g pancreatic digest of gelatin, 3g peptone, 10g lactose, 1.5g bile salt, 5g NaCl, 13.5g agar, 0.03g neutral red, 0.001g crystal violet in 1L H_2_O) and IPTG-X-gal Mueller-Hinton II agar (3g beef extract, 17.5g acid hydrolysate of casein, 1.5g starch, 13.5g agar in 1L H_2_O, supplemented with 40 mg/L X-gal and 0.2 μM IPTG) were used to differentiate LacZ(+) and (-) mutants. Cation-adjusted Mueller Hinton II Broth (MHBII) was used for antimicrobial susceptibility tests. Antibiotics were ordered from Sigma-Aldrich. Recombineering oligos were synthesized by Integrated DNA Technologies (IDT) or by the DNA Synthesis Laboratory of the Biological Research Centre (Szeged, Hungary) with standard desalting as purification.

### Oligo-mediated recombineering

Bacterial cultures *(E. coli, K. pneumoniae, C. freundii, or P. aeruginosa)* were grown in LB^L^ at 37 °C in a rotating drum. Overnight cultures were diluted 1:100, grown for 60 minutes or until OD600 ≈ 0.3, whereupon expression of SSAPs was induced for 30 minutes with 0.2% arabinose or 1 mM m-toluic acid as appropriate. Cells were then prepared for transformation. Briefly, *E. coli*, *K. pneumoniae*, and *C. freundii* cells were put on ice for approximately ten minutes, washed three times with cold water and resuspended in 1/100^th^ culture volume of cold water. This same procedure was followed for *P. aeruginosa* with the following differences: (1) Resuspension Buffer (0.5 M sucrose + 10% glycerol) was used in place of water and (2) there was no pre-incubation on ice, as competent cell prep was carried out at room temperature, which we found to be much more efficient than preparation at 4°C. After competent cell prep, 9 µl of 100 µM oligo was added to 81 µl of prepared cells for a final oligo concentration of 10 µM in the transformation mixture (2.5 µM final oligo concentration was used for *C. freundii* and *K. pneumoniae*). This mixture was transferred to an electroporation cuvette with a 0.1 cm gap and electroporated immediately on a Gene Pulser (BioRad) with the following settings: 1.8 kV (2.2 kV in the case of *P. aeruginosa*), 200 Ω, 25 µF. Cultures were recovered with SOC media for one hour and then 4 ml of LB with 1.25x selective antibiotic and 1.25x antibiotic for plasmid maintenance were added for outgrowth.

### Engineering of SEER chassis

The *E. coli* strain described in this work as the SEER chassis is engineered from EcNR2^3^. EcNR2 harbors a small piece of λ-phage integrated at the *bioAB* locus, which allows expression of λ-Red genes, and a knockout of the methyl-directed mismatch repair (MMR) gene *mutS*, which improves the efficiency of mismatch inheritance (MG1655 Δ*mutS*::*cat* Δ(*ybhB*-*bioAB*)::[*λcI857* Δ(*cro*-*orf206b*)::*tetR*-*bla*]). Modifications made to EcNR2 to engineer the SEER chassis include: 1. improvement of MAGE efficiency by mutating DNA primase (*dnaG*_Q576A)^8^, 2. introduction of a handle for SDS selection (*tolC*_STOP), 3. introduction of a handle for CHL selection (*mutS*::*cat*_STOP), and 4. removal of lambda phage with a zeocin resistance marker Δ[*λcI857* Δ(*cro*-*orf206b*)::*tetR*-*bla*]::*zeoR*. The final strain which we refer to as the SEER chassis is therefore: MG1655 Δ(*ybhB*-*bioAB*)::*zeoR* Δ*mutS*::*cat*_STOP *tolC*_STOP *dnaG*_Q576A.

### Selective allele testing in the SEER chassis

To complement the SEER chassis’ two engineered selective handles, we tested the following native antibiotic resistance alleles: [TMP: FolA P21→L, A26→G, and L28→R^62^], [KAN/GEN: 16SrRNA U1406→A and A1408→G], [SPT: 16SrRNA A1191→G and C1192→U^63^], [RIF: RpoB S512→P and D516→G^64^], [STR: RpsL K4→R and K88→R^65^], and [CIP: GyrA S83→L^66^]. 90-bp oligos conferring each mutation, with two PT bonds at their 5’ end and with complementarity to the lagging strand were designed. Two oligos were designed to repair the engineered selective handles: (1) elimination of a stop codon in the chloramphenicol acetyltransferase (*cat*) to confer CHM resistance and (2) elimination of a stop codon in *tolC* to confer SDS resistance. Oligo-mediated recombineering was run with Redβ expressed off of the pARC8 plasmid and the cultures were then plated onto a range of concentrations of the antibiotic to which the oligo was expected to confer resistance. Colony counts were made and compared to a water-blank control. Modifications targeted to provide TMP, KAN, and SPT resistance did not work adequately and so were dropped. RpsL_K43R was chosen for STR selection and RpoB_S512P for RIF selection, although in both cases there was not a significant observable difference between the two tested alleles. An antibiotic concentration was chosen that provided the largest selective advantage for those cultures transformed with oligo (Fig S2). The concentrations chosen for the selective antibiotics were: 0.1% v/v SDS, 25 μg/ml STR, 100 µg/ml RIF, 0.1 µg/ml CIP, and 20 μg/ml CHL.

### Identification of SSAP library members

To generate *Broad SSAP Library* we used a multiple sequence alignment of eight SSAPs that had been shown to function in E. coli (Redβ, EF2132 from *Enterococcus faecalis*, OrfC from *Legionella pneumophila*, S065 from *Vibrio cholerae*, Plu2935 from *Photorhabdus luminescens*, Orf48 from *Listeria monocytogenes*, Orf245 from *Lactococcus lactis*, and Bet from *Prochlorococcus siphovirus* P-SS2^19, 37^) to generate a Hidden Markov Model that described the weighted positional variance of these proteins. We then queried non-redundant nucleotide and environmental metagenomic databases using a web-based search interface^67^. Candidates were filtered based on gene size and annotation. Those that exhibited intra-sequence similarity of greater than 98% were removed from the group. We added three eukaryotic SSAP homologs to the library^68^. In total, *Broad SSAP Library* contains 120 members from the homology search, 8 members from the starting sequence alignment, and 3 eukaryotic members, or a total of 131 SSAP homologs (Table S2).

*Broad RecT Library* was generated from the full alignment of Pfam family PF03837, containing 576 sequences from Pfam 31.0^69^. Using ETE 3, a phylogenetic tree made by FastTree and accessed from the Pfam31.0 database was pruned, and from it a maximum diversity subtree of 100 members was identified (Figure S9)^70^. Five members of this group were found in Library S1, and so were excluded, and in their place six RecT variants from Streptomyces phages and eight other RecT variants were added that had previously reported activity or were otherwise of interest^6, 15, 19, 22, 23^, bringing the library size to 109 (Table S2).

### Library assembly

*Broad SSAP Library* and *Broad RecT Library* variants with a DNA barcode 22 nucleotides downstream of the stop codon were codon-optimized for *E. coli* and synthesized by Gen9 (S1) or Twist (S2). Synthesized DNA was amplified by PCR (NEB Q5 polymerase) and cloned into pDONR/Zeo (Thermo) by Gibson Assembly (NEB HiFi DNA Assembly Master Mix) and then moved into pARC8-DEST for arabinose-inducible expression. pARC8-DEST was engineered from a pARC8 plasmid^71^ that shows good inducible expression in *E. coli* by moving Gateway sites (attR1/attR2), a CHL marker, and a ccdB counter-selection marker downstream of the pBAD-araC regulatory region (Figure S1). This enabled easy, one-step cloning of the entire library into pARC8-DEST by Gateway cloning (Thermo). The Gateway reaction was transformed into E. cloni Supreme electrocompetent cells (Lucigen), providing > 10,000x coverage of both libraries in total transformants.

### Serial Enrichment for Efficient Recombineering (SEER)

Libraries were mini-prepped (NEB Monarch Kit) and electroporated into the SEER chassis with more than 1,000-fold coverage. Five cycles of oligo-mediated recombineering followed by antibiotic selection were then conducted (Fig. 1B). 5 µl of the 5 ml recovery from the recombineering step was immediately plated onto LBL + selective antibiotic plates to estimate the total throughput of the selective step. This allowed us to ensure that the library was never bottlenecked—the first round of selection was the most stringent, but we ensured that there was > 500x coverage at this stage. Following five rounds of selection, the plasmid library was mini-prepped and transformed back into the naïve parent strain, followed by five further rounds of selection (ten in total). After each selective step a 100 µl aliquot of the antibiotic-selected recovery was frozen down at −80 °C in 25% glycerol for analysis by NGS.

### Next Generation Sequencing of libraries

Primers were designed to amplify a 215 bp product containing the barcode region of the SSAP libraries from the pARC8 plasmid and to add on Illumina adaptors. PCR amplification was done with Q5 polymerase (NEB) performed on a LightCycler 96 System (Roche), with progress tracked by SYBR Green dye and amplification halted during the exponential phase. Barcoding PCR for Illumina library prep was performed as just described, but with NEBNext Multiplex Oligos for Illumina Dual Index Primers Set 1 (NEB). Barcoded amplicons were then purified with AMPure XP magnetic beads (Beckman Coulter), pooled, and the final pooled library was quantified with the NEBNext Library Quant Kit for Illumina (NEB). The pooled library was diluted to 4 nM, denatured, and a paired end read was run with a MiSeq Reagent Kit v3, 150 cycles (Illumina). Sequencing data was downloaded from Illumina, sequences were cleaned with Sickle^72^ and analyzed with custom scripts written in Python.

### Measuring recombineering efficiency in *E. coli* by NGS

To measure single locus editing, a recombineering cycle was run with an oligo that confers a single base pair non-coding mismatch in a non-essential gene (Table S4 - silent mismatch MAGE oligo 7). The allele was then amplified by PCR and editing efficiency was measured by NGS as described above. To test multiplex editing, the concentration of oligo was held fixed (10 µM in the final electroporation mixture), but the total number of oligos in the mixture was varied. Pools of oligos to test editing at 5, 10, 15, or 20 alleles simultaneously were designed so as to space the edits relatively evenly around the genome. The 5-oligo pool contained oligo #’s 3,7,11,15,17, the 10-oligo pool added oligo #’s 1,5,9,13,19, the 15-oligo pool added oligo #’s 4,8,12,16,18, and the final 20-oligo pool contained all of the silent mismatch MAGE oligos listed in Table S4. One locus (locus 8) showed major irregularities when sequenced, and so we eliminated it from our analyses.

### DIvERGE-based simultaneous mutagenesis of *gyrA, gyrB, parE,* and *parC*

A single round of DIvERGE mutagenesis was carried out to simultaneously mutagenize *gyrA, gyrB, parE*, and *parC* in *E. coli* MG1655 by the transformation of an equimolar mixture of 130 soft-randomized DIvERGE oligos, tiling the four target genes. The sequences and composition of these oligos were published previously (Nyerges, A., et. al, PNAS, 2018)^13^. To perform DIvERGE, 4 µl of this 100 µM oligo mixture was electroporated into *E. coli* K-12 MG1655 cells expressing Redβ from pORTMAGE311B, PapRecT from pORTMAGE312B, or CspRecT from pORTMAGE-Ec1, in 5 parallel replicates according to our previously described protocol^46^. Following electroporation, the replicates were combined into 10 ml fresh TB media. Following recovery for 2 hours, cells were diluted by the addition of 10 ml LB and allowed to reach stationary phase at 37 °C, 250 rpm.

Library generation experiments were performed in triplicates. Following library generation, 1 mL of outgrowth from each library was subjected to 250, 500, and 1,000 ng/mL ciprofloxacin (CIP) stresses on 145 mm-diameter LB-agar plates. Colony counts were determined after 72 hours of incubation at 37 °C, and individual colonies were subjected to further genotypic (i.e., capillary DNA sequencing) analysis and phenotypic (i.e., Minimum Inhibitory Concentration) measurements.

### pORTMAGE plasmid construction and optimization

All plasmids used in the study are listed in Table S7. Cloning reactions were performed with Q5 High-Fidelity Master Mix and HiFi DNA Assembly Master Mix (New England Biolabs). pORTMAGE312B (Addgene accession: 128969) and pORTMAGE-Ec1 (Addgene accession: 138474) were constructed by replacing the Redβ open reading frame (ORF) of pORTMAGE311B plasmid (Addgene accession: 120418)^43^ with PapRecT and CspRecT respectively. pORTMAGE-Pa1 was constructed in many steps: i.) the Kanamycin resistance cassette and the RSF1010 origin-of-replication on pORTMAGE312B with Gentamicin resistance marker and pBBR1 origin-of-replication, amplified from pSEVA631^73^ (gift from Prof. Victor de Lorenzo - CSIC, Spain), ii.) optimization of RBSs in pORTMAGE-Pa1 was done by designing a 30-nt optimal RBS in front of the SSAP ORF and in between the SSAP and MutL ORFs with an automated design program, De Novo DNA^74^, iii.) *Pa*MutL was amplified from *Pseudomonas aeruginosa* genomic DNA and cloned in place of *Ec*MutL_E32K, and finally iv.) *Pa*MutL was mutated by site-directed mutagenesis to encode E36K. Ssr and Rec2 were ordered as gblocks from IDT and cloned in place of PapRecT into earlier versions of pORTMAGE-Pa1 for the comparisons in Figure S6.

### Measuring recombineering efficiency in Gammaproteobacteria by selective plating

Oligos were designed to introduce I) premature STOP codons into *lacZ* for *E. coli*, *K. pneumoniae*, and *C. freundii*, or II) RpsL K43→R; GyrA T83→I; ParC S83→L; RpoB D521→V, or a premature STOP codon into *nfxB* for *P. aeruginosa*. Oligo-mediated recombineering was performed as described above on all bacterial strains. After recovery overnight, cells were plated at empirically-determined dilutions to a density of 200-500 colonies per plate. In the case of LacZ screening, plating was assayed on MacConkey agar plates or on X-Gal/IPTG LB^L^ agar plates in the case of *K. pneumoniae*. In the case of selective antibiotic screening, cultures were plated onto both selective and non-selective plates. Selective antibiotic concentrations used were the same as those described for the selective testing above, except that in *P. aeruginosa* 100 µg/ml STR and 1.5 µg/ml CIP were used unless otherwise noted. Variants that were resistant to multiple antibiotics were selected on LB^L^ agar plates that contained the combination of all corresponding antibiotics. Non-selective plates were antibiotic-free LB^L^ agar plates. In all cases, allelic-replacement frequencies were calculated by dividing the number of recombinant CFUs by the number of total CFUs. Plasmid maintenance was ensured by supplementing all media and agar plates with either KAN (50 µg/ml) or GEN (20 µg/ml).

### Minimum Inhibitory Concentration (MIC) measurement in *P. aeruginosa*

MICs were determined using a standard serial broth microdilution technique according to the CLSI guidelines (ISO 20776-1:2006, Part 1: Reference method for testing the in vitro activity of antimicrobial agents against rapidly growing aerobic bacteria involved in infectious diseases). Briefly, bacterial strains were inoculated from frozen cultures onto MHBII agar plates and were grown overnight at 37 °C. Next, independent colonies from each strain were inoculated into 1 ml MHBII medium and were propagated at 37 °C, 250 rpm overnight. To perform MIC tests, 12-step serial dilutions using 2-fold dilution-steps of the given antibiotic were generated in 96-well microtiter plates (Sarstedt 96-well microtest plate). Antibiotics were diluted in 100 μl of MHBII medium. Following dilutions, each well was seeded with an inoculum of 5×10^4^ bacterial cells. Each measurement was performed in 3 parallel replicates. Plates were incubated at 37 °C under continuous shaking at 150 rpm for 18 hours in an INFORS HT shaker. After incubation, the OD_600_ of each well was measured using a Biotek Synergy 2 microplate reader. MIC was defined as the antibiotic concentration which inhibited the growth of the bacterial culture, i.e., the drug concentration where the average OD600 increment of the three replicates was below 0.05.

## Acknowledgements

We thank Erkin Kuru, Max Schubert, Aditya Kunjapur, and Devon Stork for helpful discussions, and Prof. Victor de Lorenzo (Centro Nacional de Biotecnologia, CSIC, Spain) for providing plasmids for the study. Claire O’Callaghan, and Verena Volf provided experimental help in various capacities in closely related work. John Aach was helpful for thinking through conceptual ideas. Funding for this research was graciously provided by the DOE under grant DE-FG02-02ER63445 (to G.M.C), the European Research Council H2020-ERC-2014-CoG 648364 ‘Resistance Evolution’ (to C.P.), GINOP (MolMedEx TUMORDNS) GINOP-2.3.2-15-2016-00020, GINOP (EVOMER) GINOP-2.3.2-15-2016-00014 (to C.P.); the ‘Lendület’ Program of the Hungarian Academy of Sciences (to C.P.), and a PhD fellowship from the Boehringer Ingelheim Fonds and an EMBO Long-Term Fellowship (to Á.N.). M.C. was supported by the Szeged Scientists Academy under the sponsorship of the Hungarian Ministry of Human Capacities (EMMI: 13725-2/2018/INTFIN), by the UNKP-18-2 New National Excellence Program of the Hungarian Ministry of Human Capacities and UNKP 10-2 New National Excellence Program of the Hungarian Ministry for Innovation and Technology.

## Supplementary Figures

**Figure S1.**
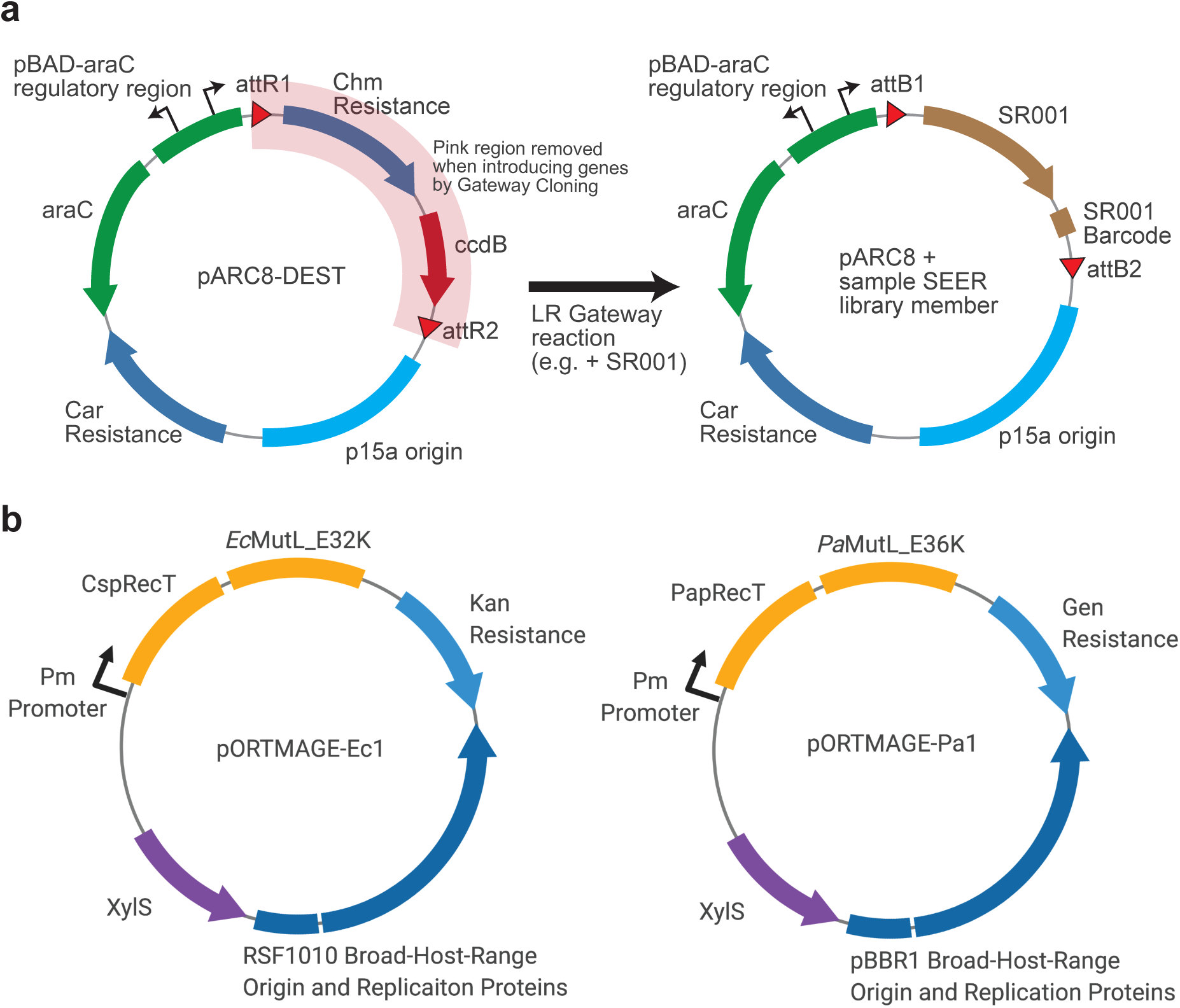
Vector maps. (A) pARC8-DEST was created to have a pBAD regulatory region, beta lactamase, a p15a origin, and a lethal *ccdB* gene flanked by attR sites for Gateway cloning. Introduction by the LR Gateway reaction of for instance SR001, would create the vector on the right, with an arabinose-inducible SR001 followed by a barcode. (B) Two pORTMAGE vectors are provided for broad-spectrum recombineering. pORTMAGE-Ec1 was demonstrated effective in *E. coli*, *C. freundii*, and *K. pneumoniae*, while pORTMAGE-Pa1 was demonstrated effective in *P. aeruginosa*.

**Figure S2.**
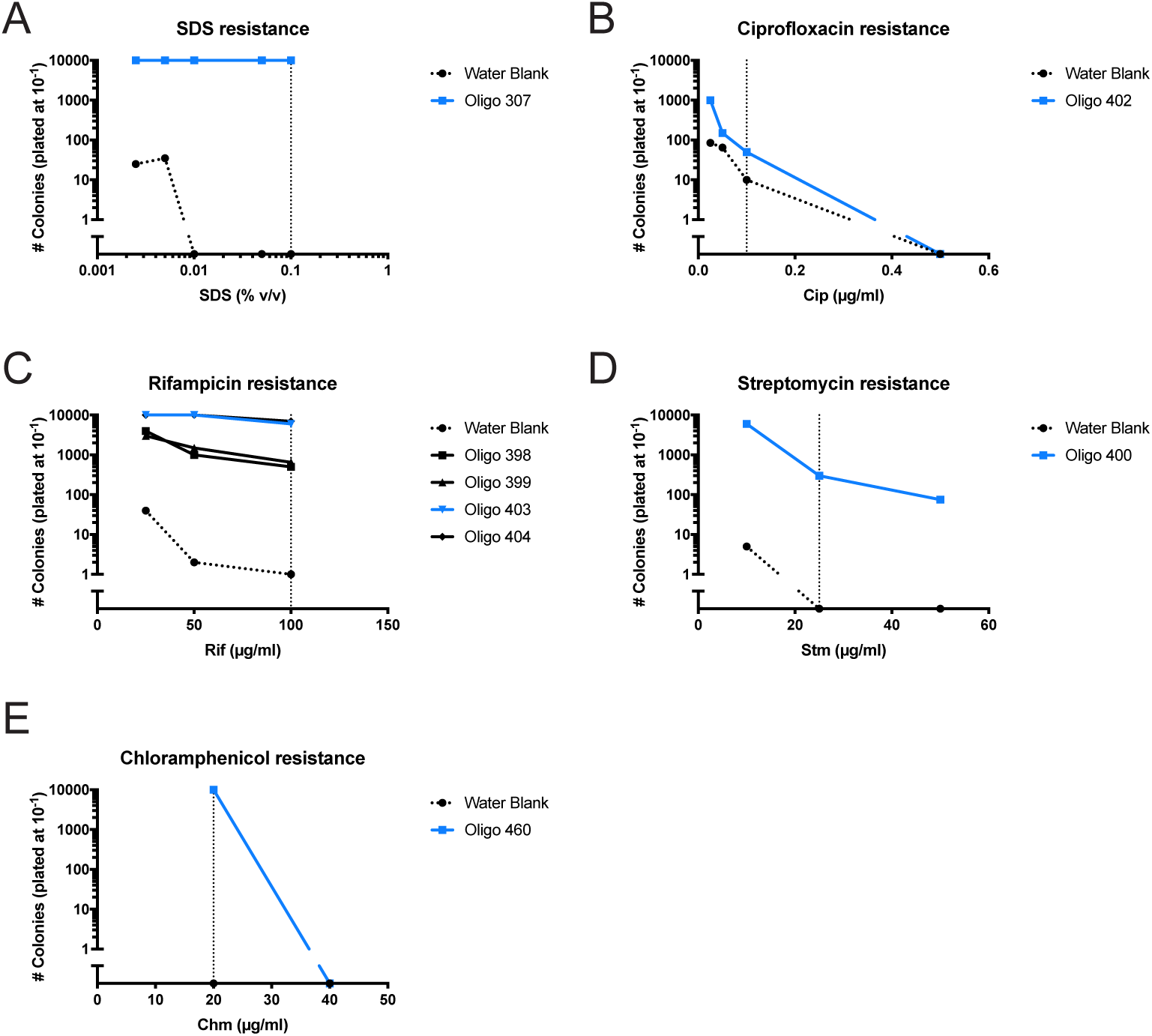
Antibiotic testing in *E. coli*. Candidate oligos used to confer resistance to the five antibiotic selections used in the SEER chassis were tested for their efficacy at varied concentrations of antibiotic. The number of CFUs is plotted against antibiotic concentration for each antibiotic. Oligos that were used in SEER selection are colored light blue. The antibiotic concentration chosen for the SEER selection is indicated by a vertical dashed line. (A) SDS resistance was restored by correcting a stop codon insertion in *tolC*. (B) Ciprofloxacin, (C) Rifampicin, and (D) Streptomycin resistance were imparted with a mutation to native *E. coli* genes. (E) Chloramphenicol resistance was restored by correcting a stop codon insertion in *cat*.

**Figure S3.**
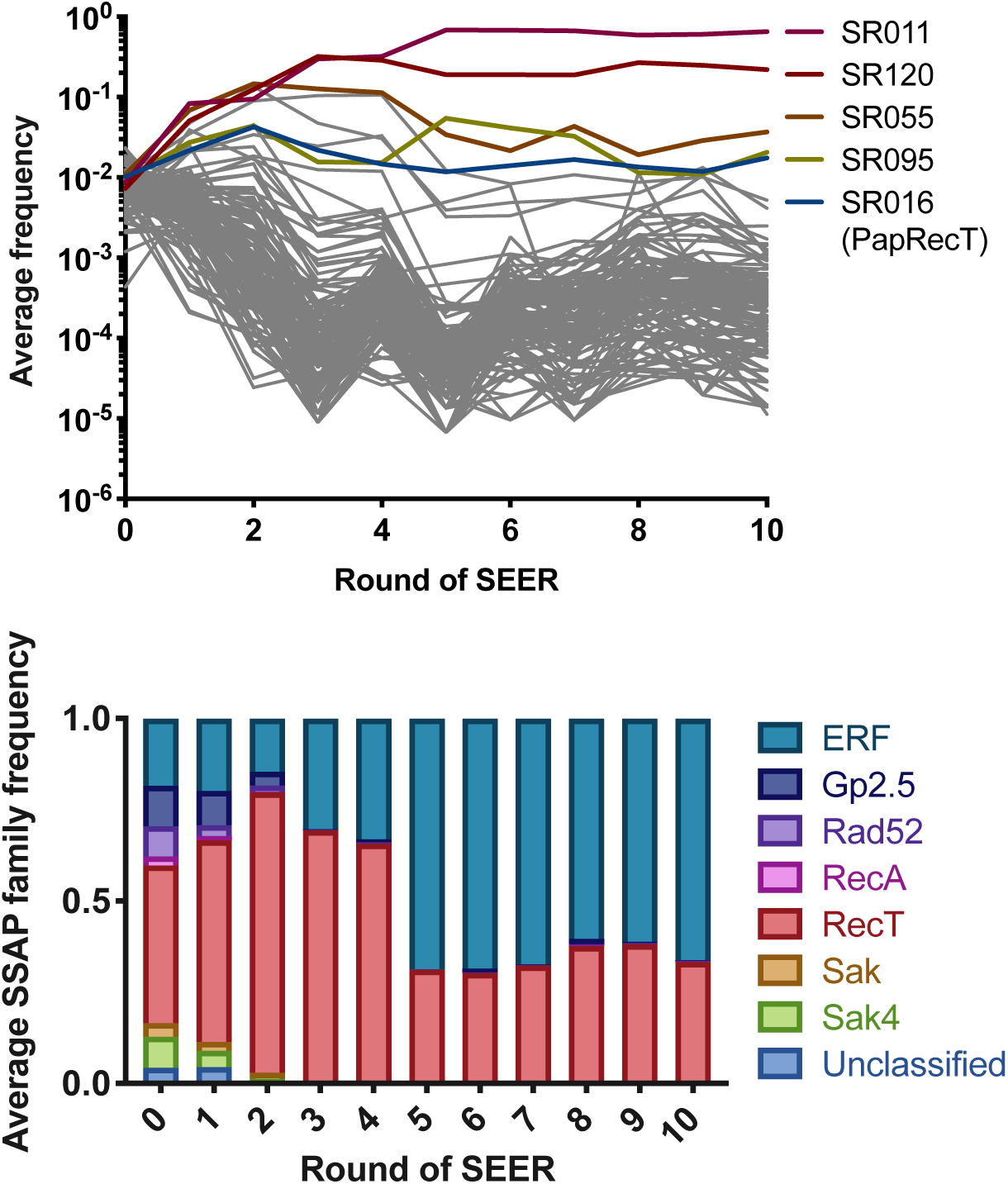
(Top) the frequency of each library member from the *Broad SSAP Library* is plotted across ten rounds of SEER. Data are measured by amplification of the barcoded region of the plasmid library and next generation sequencing (NGS). (Bottom) The frequency of SSAP families is shown across the same selective cycles on the *Broad SSAP Library*.

**Figure S4.**
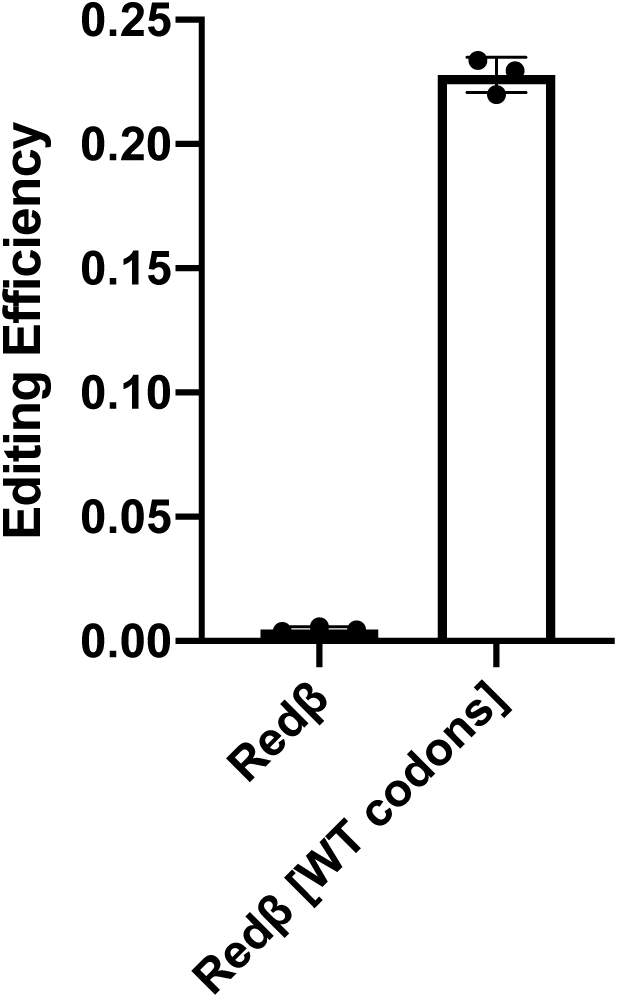
The effect of codon-usage on Redβ editing efficiency in *E. coli*. The efficiency of Redβ from the *Broad SSAP Library* was compared with Redβ expressed off of its wild-type codons. Efficiency of making a single base pair mutation in a non-coding gene (silent mismatch MAGE oligo 7) was measured by NGS.

**Figure S5.**
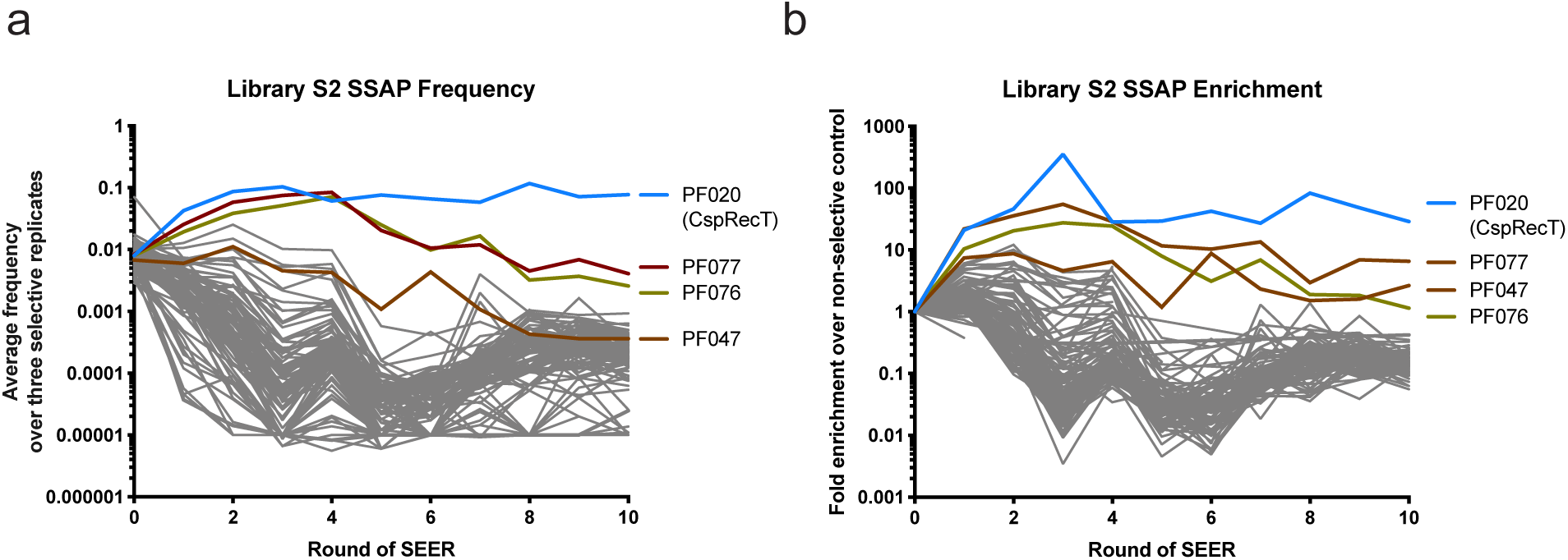
Frequency and Enrichment of members of *Broad RecT Library* over ten rounds of SEER enrichment.

**Figure S6.**
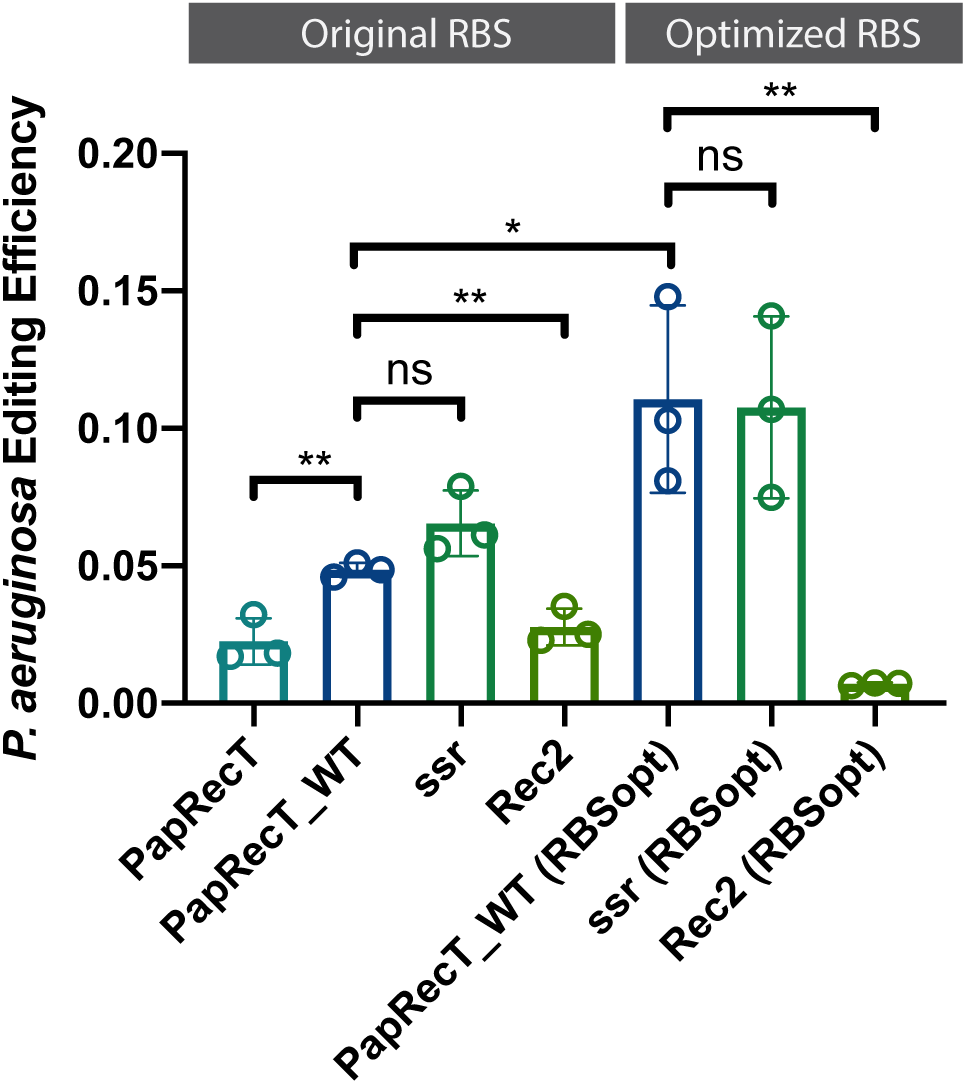
Recombineering efficiency in *P. aeruginosa* was measured for PapRecT with *E. coli* codons, PapRecT with its wild-type codons, and two SSAPs that have been reported to work in *Pseudomonas putida*. This was measured both with the original pORTMAGE311B RBS and an RBS optimized for *P. aeruginosa*. Significance values are indicated for a parametric t-test between two groups, where ns, *, **, ***, and ***** indicate p > 0.05, p < 0.05, p < 0.01, p < 0.001, and p < 0.0001 respectively.

**Figure S7.**
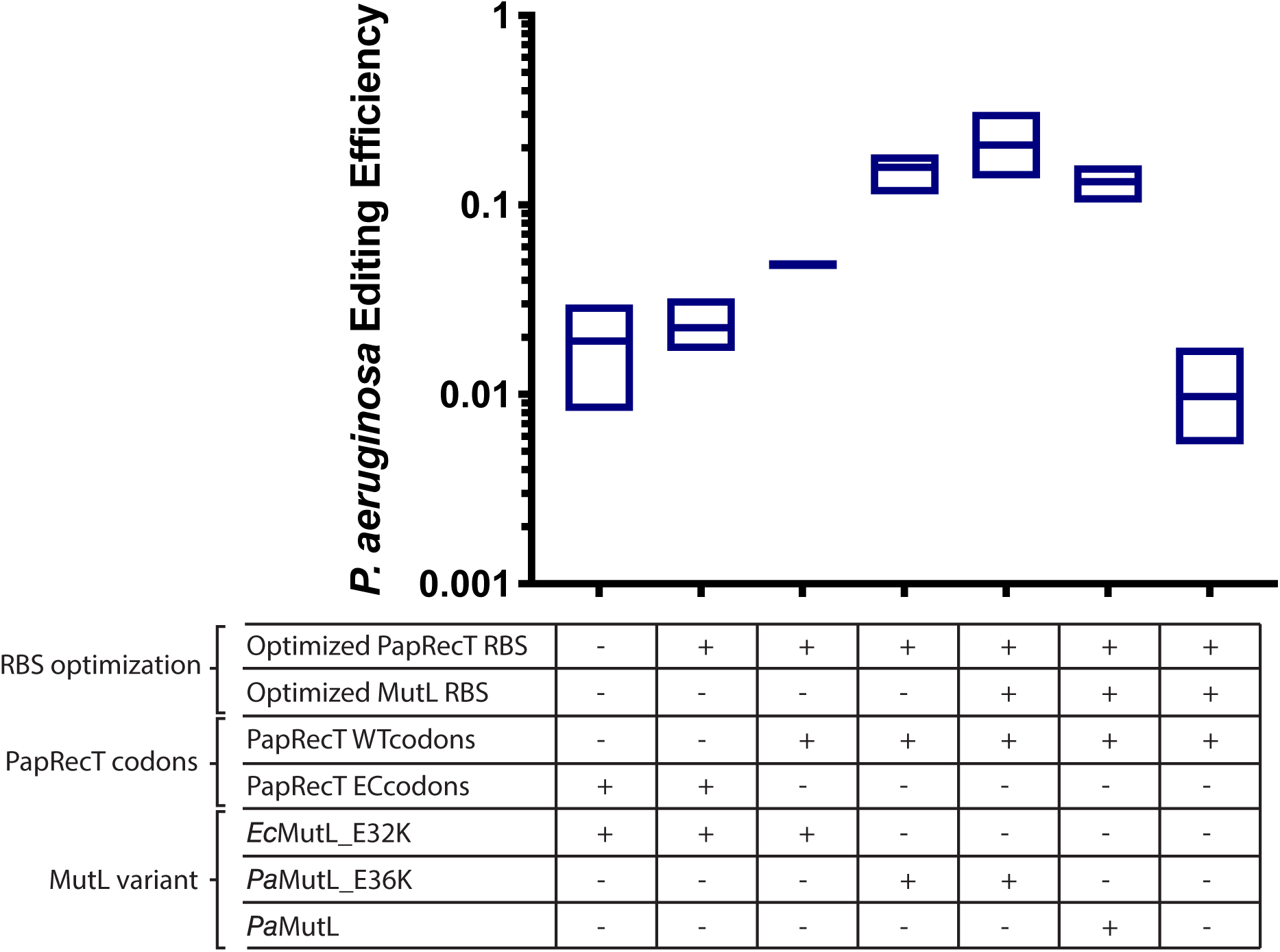
Editing efficiency in making a single-base mutation at the *rpsL* locus in *P. aeruginosa* with various plasmid variants expressing PapRecT. An unoptimized plasmid (far left) was constructed by replacing, in pORTMAGE312B (Addgene accession: 128969), the RSF1010 origin of replication and the kanamycin resistance gene with a pBBR1 origin of replication and a gentamicin resistance gene. The best-performing plasmid variant (third from right) was renamed pORTMAGE-Pa1 (Addgene accession: 138475). Constructs examining the role of MutL in single-base recombineering efficiency were made by first restoring wild-type *Pa*MutL and then by removing it entirely (second from right, and far right respectively).

**Figure S8.**
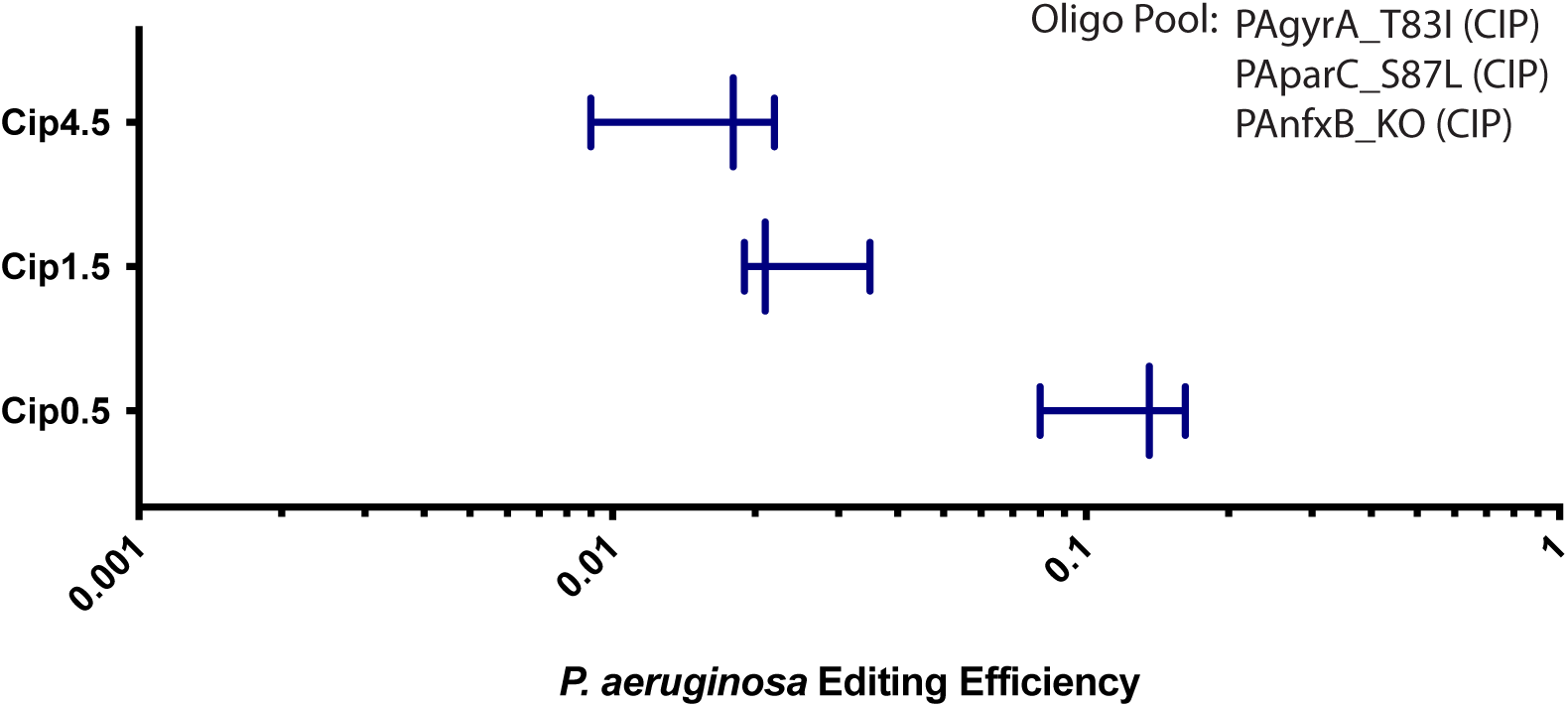
One round of MAGE with a pool of three oligos that confer Ciprofloxacin resistance was conducted in *P. aeruginosa* with pORTMAGE-Pa1. Editing efficiency is shown after plating on three different concentrations of antibiotic.

**Figure S9.**
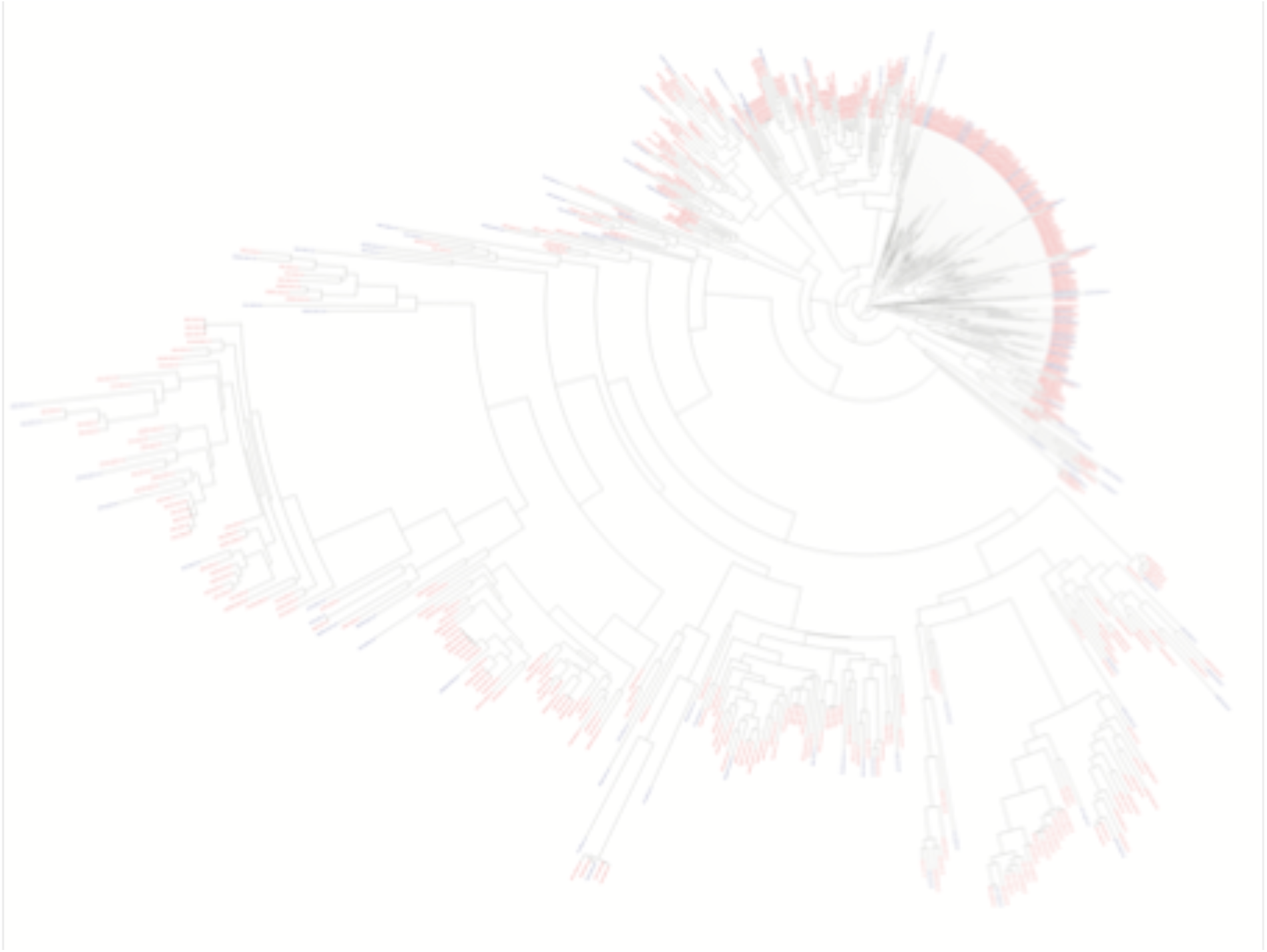
Phylogenetic tree of the Pfam RecT protein family. The *Broad RecT Library* was generated from the full alignment of Pfam family PF03837, containing 576 sequences from Pfam 31.0^69^, as generated by FastTree. Using ETE 3, the phylogenetic tree was pruned and a maximum diversity subtree of 100 members was identified^70^. Members of this 100-member subtree are indicated in blue.

**Figure S10.**
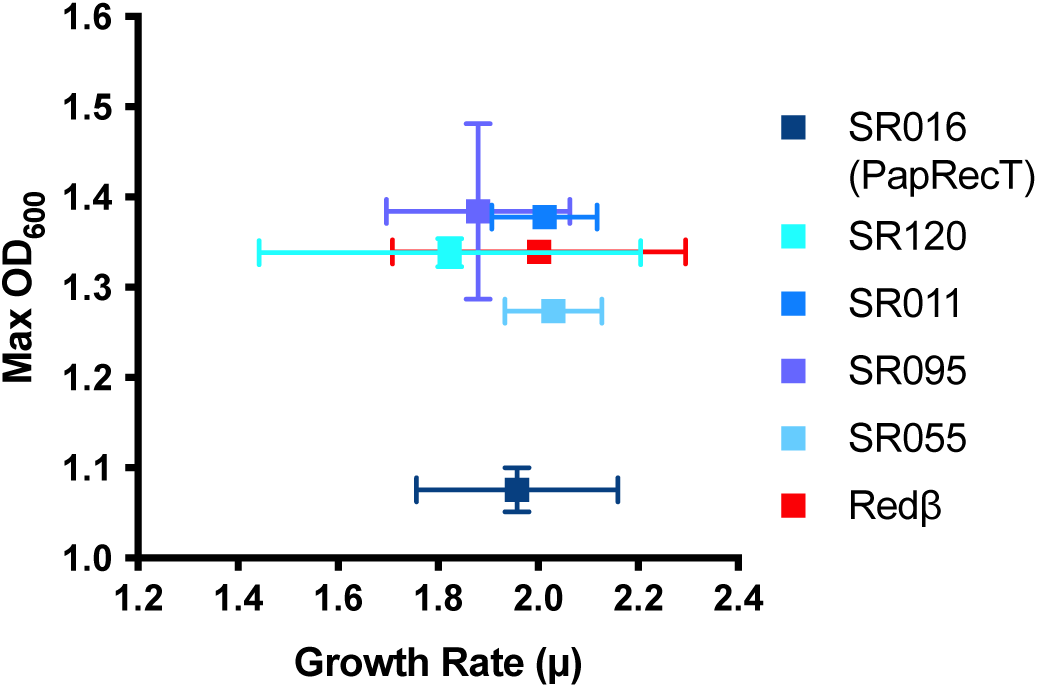
The growth rates of bacteria expressing each of the top five candidates from *Broad SSAP Library* or Redβ were measured by plate-reader growth assay and plotted against the maximum attained OD_600_ of the culture.

